# Inhibition benefits neural system identification

**DOI:** 10.64898/2026.05.27.728086

**Authors:** Yuyao Deng, Zhuokun Ding, Jiakun Fu, Jonathan Oesterle, Andreas S. Tolias, Thomas Euler, Yongrong Qiu

**Affiliations:** Institute for Ophthalmic Research, University of Tübingen, Tübingen, Germany; Graduate Training Centre of Neuroscience, University of Tübingen, Tübingen, Germany; Department of Ophthalmology, Byers Eye Institute, Stanford University School of Medicine, Stanford, CA, USA; Stanford Bio-X, Stanford University, Stanford, CA, USA; Wu Tsai Neurosciences Institute, Stanford University, Stanford, CA, USA; Salk Institute for Biological Studies, CA, USA; Hertie Institute for AI in Brain Health, University of Tübingen, Tübingen, Germany; Department of Electrical Engineering, Stanford University, Stanford, CA, USA; Bernstein Centre for Computational Neuroscience, Tübingen, Germany; Centre for Integrative Neuroscience, University of Tübingen, Tübingen, Germany

## Abstract

*Neural system identification* approaches use empirical data to fit the stimulus-response functions of neurons. Augmented by deep neural networks, such models have achieved high predictive performance and allow to perform *in silico* experiments to test hypotheses. Yet, many of these methods ignore common features in visual systems, such as inhibitory interactions between neurons, which are essential for nonlinear neural computation. Here, we incorporate inhibition as an inductive bias into a deep model for neural prediction and investigate the influence of inhibition on the learned transfer functions. To this end, we employ difference-of-Gaussian (subtraction) and within-channel divisive normalization (division), which have been proposed to relate inhibition to neural processing, in deep networks for predicting visual responses. We observe that incorporating such operations maintains the predictive performance and encourages the learning of biologically plausible kernels reminiscent of neural representation in early vision. Additionally, our *in silico* experiments demonstrate that implementing either subtractive or divisive operation benefits the learning of surround suppression but not cross-orientation inhibition. Interestingly, while division increases the sparsity of activation and reduces the sparsity of weights, subtraction has the reverse effect.

## 1 Introduction

A primary task of computational neuroscience is to predict neuronal responses to diverse stimuli. This endeavor has been advanced through *neural system identification* methods, which learn transfer functions by mapping stimuli to responses based on empirical data [Wu et al., 2006]. Classical approaches, such as linear-nonlinear Poisson models and subunit models, aim to capture the nonlinear processing inherent in visual circuits [e.g., Chichilnisky, 2001, Rust et al., 2005, Duncker et al., 2023, Sridhar et al., 2024]. More recently, inspired by hierarchical computation in neural systems, deep neural networks (DNNs) have emerged as powerful tools for response prediction. In particular, within visual systems, DNN models have demonstrated remarkable efficacy in identifying stimulus-response functions and testing new hypotheses with *in silico* experiments, encompassing regions such as the retina [Batty et al., 2016, McIntosh et al., 2016, Hoefling et al., 2022] and visual cortex [Ponce et al., 2019, Bashivan et al., 2019, Wu et al., 2024, Tong et al., 2023, Turishcheva et al., 2024a].

Such data-driven DNNs were initially inspired by the brain, yet many of them do not consider inhibitory interactions between neurons, which are key in biological visual systems [Barlow, 1953, Hartline et al., 1956]. These interactions are crucial to efficient neural computation and adaptive information processing, contributing to processes like surround modulation and spatial frequency tuning. Many computational models have been proposed to explain how inhibition might regulate neuronal responses. For example, the Difference-of-Gaussian (DoG) model use lateral inhibition to represent center-surround interactions by subtracting surround-induced activity [Rodieck, 1965, Enroth-Cugell and Robson, 1966]. Divisive normalization (DN), on the other hand, ensures balanced processing by dividing each neuron’s activity by the sum of responses of nearby neurons [Albrecht and Geisler, 1991, Heeger, 1992, Carandini and Heeger, 2012]. These operations, subtraction and division, though simple, effectively simulate physiological experimental observations including surround suppression [Cavanaugh et al., 2002a,b], cross-orientation inhibition [Morrone et al., 1982, Freeman et al., 2002], and multisensory integration [Ohshiro et al., 2011].

Inhibition has been integrated into classical methods for neural prediction. For example, Burg and colleagues incorporated canonical divisive normalization - defined as normalization by pooled activity across feature channels - into a subunit model and showed that this constraint enabled the emergence of cross-orientation inhibition without degrading predictive performance. This work demonstrates that explicitly modeling inhibitory computations, such as canonical divisive normalization [Albrecht and Geisler, 1991, Carandini and Heeger, 2012], can shape response properties in subunit models. We then ask whether deep networks with high predictive performance can benefit from inhibition to recover functional features.

Here, we incorporate inhibitory computations as an inductive bias into a DNN model for the identification of response functions, focusing on the effect of interactions between neurons by means of subtraction and division on learned neural features (Figure 1). We decompose each feature map into spatially defined center and surround components and recombine them using either a subtractive or a divisive operation. The subtractive operation was inspired by classical Difference-of-Gaussians (DoG) models, whereas the divisive operation was derived from the framework of divisive normalization (DN) described by Matteo Carandini and David J. Heeger[Carandini and Heeger, 2012]. In our work, both subtraction and division are applied to spatially antagonistic signals within a feature map. To this end, we build a DNN to estimate the stimulus-response functions of neurons in the mouse visual cortex. We find that our models incorporating inhibition – by using subtractive and divisive operations – have equivalent or better predictive performance compared to those without inhibition. Notably, subtraction and division encourage networks to learn biologically similar kernels reminiscent of neural representation in early vision. Next, our *in silico* experiments demonstrate that subtractive and divisive operations benefit the learning of surround suppression, but not cross-orientation. Finally, our simulation suggests that division reduces connectivity sparsity and improves activation sparsity, while subtraction has inverse effects.

**Figure 1:**
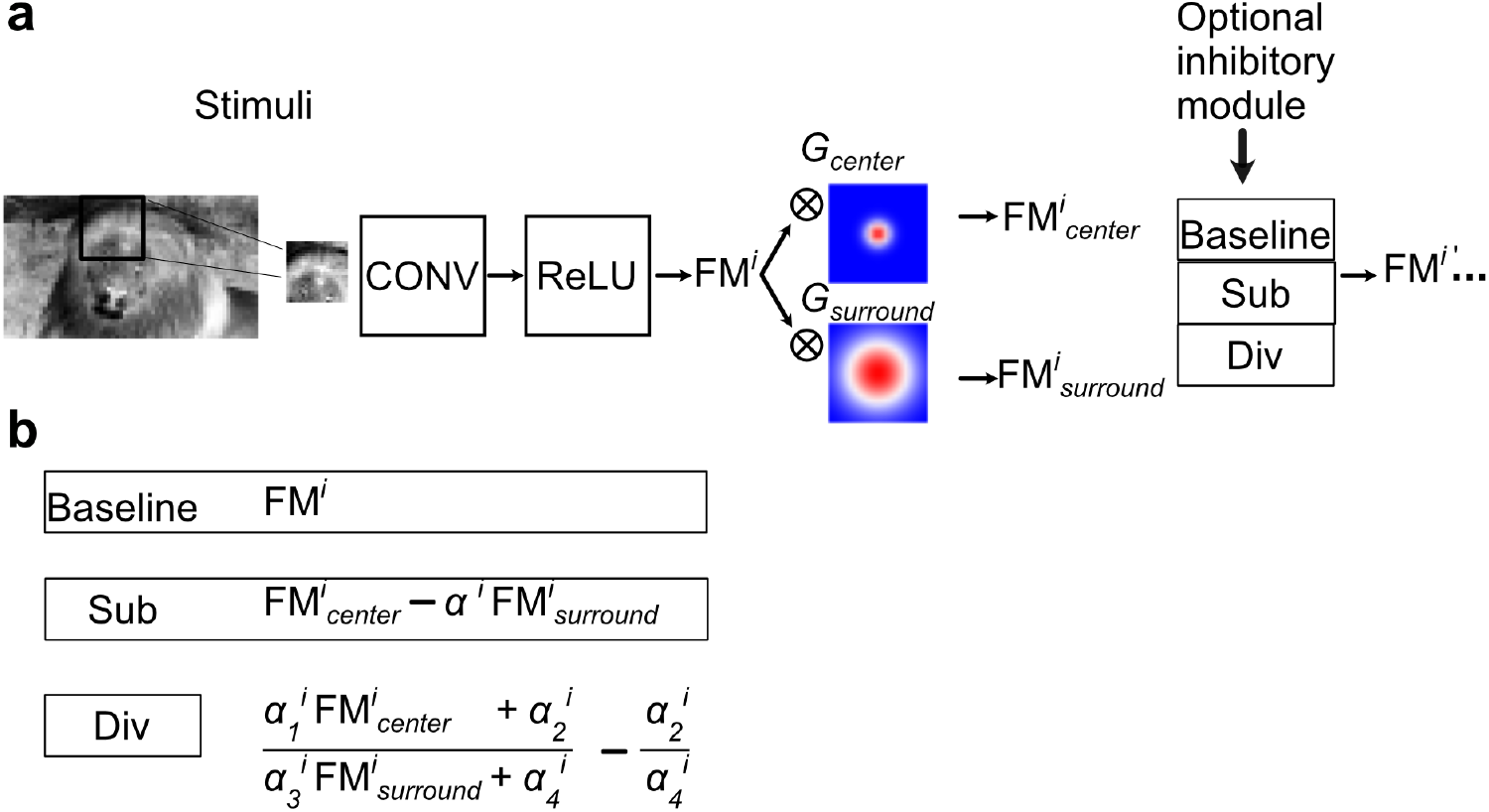
Illustration of integrating inhibitory processing into a DNN model for system identification. Deep networks comprise of linear and nonlinear computations such as the convolution layer and the rectified linear unit (ReLU) layer, aiming at predicting responses to visual stimuli. Our approach explicitly implements inhibition between nearby neurons by convolving each feature map (*FM* ^*i*^) with 2D Gaussians (*G*_*center*_ and *G*_*surround*_) and applying subtractive or divisive operations on the yielded features (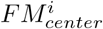 and 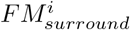).

### 2 Materials and methods

### 2.1 Dataset

We used recordings from two animals from a publicly available dataset [Franke et al., 2022], which consists of light-evoked responses from neurons in the mouse primary visual cortex (V1) to static dichromatic natural images (UV/green, 36 × 64 pixels), recorded with two-photon Ca^2+^ imaging. To quantify how reliably a cell responded to a stimulus, we calculated, as a proxy for the signal-to-noise ratio, a quality index (*QI*):

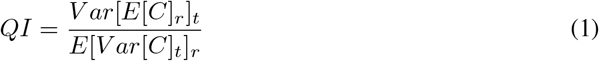

where *C* is the response matrix with a shape of *t × r* (time samples by repetitions of the stimulus). *E*[]_*x*_ and *V ar*[]_*x*_ denote the mean and variance in the indicated dimension *x*, respectively. After applying a response quality filter to 10-repeat test responses with *QI >* 0.3, we obtained 161 neurons and 91 neurons from each animal scan respectively for modeling. The single-trial responses of these neurons to 4,400 images were split into 4,000 for training and 400 for validation. The mean responses across ten repetitions to 79 test images were used to evaluate model performance.

### 2.2 Inhibition models

To integrate inhibitory processing into the DNN model, inspired by previous work on implementing subtractive or divisive normalization for image recognition [Hasani et al., 2019, Miller et al., 2021, Pogoncheff et al., 2023] or for modeling responses to parametric stimuli [Aqil et al., 2021], we convolved each feature map (*FM* ^*i*^, *i* representing channel index) with two 2D Gaussian kernels (*G*_*center*_ and *G*_*surround*_; Figure 1). These two kernels were predefined with different standard deviations, with *G*_*center*_’s standard deviation being smaller than *G*_*surround*_’s. Each Gaussian kernel was defined as follows:

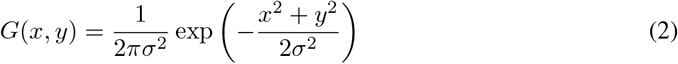

where *x* and *y* represent the pixel positions relative to the center of the kernel, and *σ* is the standard deviation. We normalized the kernel so that its elements sum to 1. Practically, we used a convolutional kernel 1 × 1 for *G*_*center*_, yielding an output 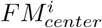 being the same as the input *FM* ^*i*^.

We employed *G*_*surround*_ with a size of 5 × 5 and a standard deviation *σ* = 1.0, generating an output 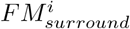 being smoother than *FM* ^*i*^. We next implemented inhibitory interactions on the generated 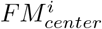 and 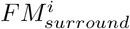 using either subtraction or division.

#### Subtractive module

The subtractive operation was defined as:

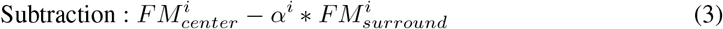

Here, each channel is regulated by a learnable parameter *α*_*i*_, which was initialized with 1.0.

#### Divisive module

The divisive operation was defined as

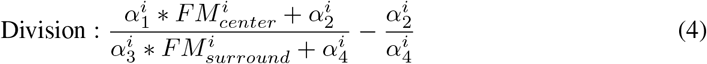

where four learnable parameters 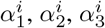 and 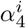 were initialized with 1.0, 0.0, 1.0 and 10.0. All aforementioned initialization values were kept unchanged when training models on different animals.

### 2.3 Baseline and control models

We used a DNN without normalization as our baseline model (Figure 1), applying L2 regularization to the weights of convolutional layers and L1 regularization to the weights of fully connected layers. Specifically, the baseline model consisted of a convolutional layer (48 × 2 × 9 × 9, output channels × input channels × kernel width × kernel height), followed by a ReLU function, a second convolutional layer (48 × 48 × 7 × 7) and another ReLU function. The resulting feature maps were flattened and fed into a fully connected layer, followed by an exponential activation function. We used a stride of 1 and no padding for convolutional layers. We trained each model on the training dataset, optimized its hyperparameters using the validation data, and evaluated it on the test data.

To investigate the effects of inhibition, we implemented a subtraction (Sub) or division (Div) module immediately after the first rectifying function (ReLU1). For control models, we added either a batch normalization layer (`torch.nn.BatchNorm2d`), or a layer normalization (`torch.nn.LayerNorm`), or a group normalization layer (`torch.nn.GroupNorm`) immediately after ReLU1 in the baseline network, which constituted BN, LN, and GN models, respectively. Based on these three control models, we added subtractive or divisive modules after their respective normalization layer, generating their inhibitory model variants: BN_Sub, BN_Div, LN_Sub, LN_Div, GN_Sub, GN_Div. All 12 models’ architecture are sketched in Suppl. Figure A.1. We trained all 12 models on one animal (n = 161 cells) and Baseline, Sub, and Div on another animal (n=91).

### 2.4 Training and analysis

#### Model training

We trained all models with a learning rate of 2 × 10^−4^ for up to 200 epochs using the Adam optimizer. For the parameter *α*_*i*_ in the subtraction model, we used a learning rate of 0.02.

The loss function was defined as:

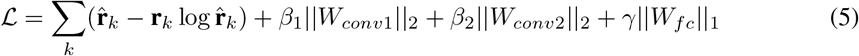

Here, the first term represents the Poisson loss between predicted responses 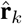 and recorded responses **r**_*k*_ with neuron index *k*, the second and third terms represent L2 penalties on the weights of convolutional filters (*W*_*conv*1_ and *W*_*conv*2_) with hyperparameters *β*_1_ and *β*_2_, and the fourth term represents the L1 penalty on the weights of the fully connected layer with hyperparameter *γ*. We quantified the predictive performance of each model by computing the Pearson correlation coefficient (CC) between the predicted and recorded responses. We tuned the hyperparameters *β*_1_, *β*_2_ and *γ* through a grid search on the validation data and selected the configuration that achieves the best performance. Additionally, we computed the log-likelihood on the test data as another metric to evaluate the models. For both metrics, higher values indicate better model performance.

#### Biological plausibility

To quantify how the learned feature maps resembled biologically plausible receptive fields, we fitted three parametric functions to each kernel from the first convolutional layer (48 × 2 × 9 × 9): a two-dimensional Gaussian, a Difference of Gaussian, and a two-dimensional Gabor function. We used `scipy.optimize.curve_fit` and computed the coefficient of determination (*R*^2^) to quantify the goodness of fit as follows:

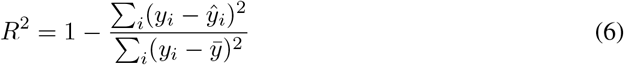

where *y*_*i*_ represents the original filter values, *ŷ*_*i*_ represents the fitted values, and 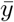 is the mean of the original filter values. For each trained model, this procedure produced 96 (48 feature maps per color channel) *R*^2^ per parametric fitting function. We used the highest *R*^2^ value among the three functions for each kernel as a proxy of biological plausibility. A higher *R*^2^ indicates that the filter can be well approximated by one of the canonical receptive field shapes.

#### Optimal Gabor identification

We first identified the optimal Gabor that elicited the highest response for each neuron by performing *in silico* experiments on trained models. The Gabor filter was defined as:

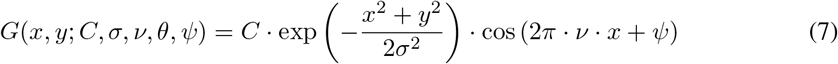

where *x* and *y* denote the spatial coordinates after centering and rotation. These two procedures are defined as:

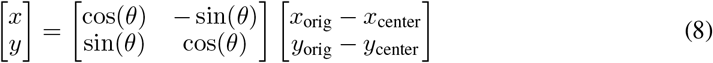

Here, *x*_orig_ and *y*_orig_ are the spatial grid coordinates before rotation, *x*_center_ and *y*_center_ are the center of the Gabor patch, *C* denotes the contrast, *σ* the standard deviation of the Gaussian envelope. *θ, ν* and *ψ* represent the orientation, the spatial frequency, and the phase shift of the sinusoidal grating. We systematically varied these parameters to determine the Gabor that maximizes each neuron’s activity for each trained model: We sampled (*x*_0_, *y*_0_) from a grid with 2-pixel steps across the 36 × 64 space, where *x*_0_ = 0, 2, 4, …, 62 and *y*_0_ = 0, 2, 4, …, 34. We shifted *C* with equidistant values in log space *C* = 0.01 *× C*_*max*_ × (2.51189)^*i*^ for *i* = 0, 1, …, 5, where *C*_*max*_ represents the maximum contrast level allowed, ensuring that the generated Gabor contrast values remain within the valid range of the normalized training data. We varied the orientations of the sinusoidal grating at evenly spaced angles *θ* = 0, 1/12, 2/12, …, 11/12 (in units of *π*), its phases at *ψ* = 0, 1/8, 2/8, …, 7/8 (in units of 2*π*) and its spatial frequency according to *ν* = 1/1.3 × (1.3)^*i*^ for *i* = 0, 1, …, 9. We ranged the size of the Gaussian envelope by setting the standard deviation *σ* = 0.9 × (1.3895)^*i*^ for *i* = 0, 1, …, 7, where the maximum *σ* was 9 pixels. Suppl. Figures A.6 illustrates the dimensions of Gabors we manipulate during the search for the optimum.

#### Suppression index

After identifying the optimal Gabor filter for each neuron, we estimated the size-tuning curve for each neuron per model. We defined the suppression index (*SI*) as [Cavanaugh et al., 2002a]:

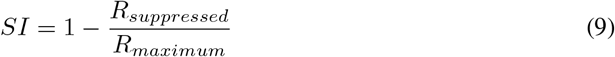

where *R*_*maximum*_ denotes the maximal activity driven by the Gabor stimulus set with varying sizes and *R*_*suppressed*_ represents the response to the Gabor stimulus with the largest standard deviation. The higher *SI* indicates stronger surround inhibition. We use *SI >* 0.1 as threshold to select neurons that exhibit surround suppression.

#### Cross-orientation inhibition index

With the optimal Gabor obtained, we generated its orthogonal Gabor by rotating the optimal one by 90°. After acquiring the optimal and orthogonal Gabor pair for all model neurons, we superimposed the orthogonal Gabor onto the optimal one, varying both Gabors independently across nine contrast levels, which produced a 9 × 9 stimulus grid. We fed the superimposed stimulus to the model and computed a response matrix *R*_*grid*_(*i, j*) (*i*, optimal Gabor contrast; *j*, orthogonal Gabor contrast) for each neuron. We then calculated the cross-orientation inhibition (*COI*) index as follows:

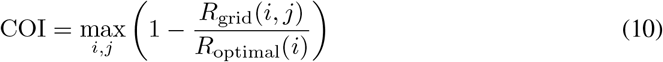

where *R*_*optimal*_(*i*) indicates the response to the optimal Gabor at the contrast level *i* without the superimposed orthogonal Gabor. A value of *COI* close to one indicates strong suppression by the orthogonal Gabor component, while a value near zero indicates weak or no suppression. We defined a neuron as cross-orientation suppressive if its response diminishes by more than 10% in the presence of an orthogonal Gabor filter [Burg et al., 2021].

#### Weight and activation sparsity

We quantified sparsity in both model weights and activations using three complementary metrics: Kurtosis, the Gini coefficient, and Hoyer sparsity. Weight sparsity was computed from the first convolutional layer (Conv1), which captures unit-to-unit connectivity. Activation sparsity was analyzed at two levels: Lifetime sparsity measures how frequently an individual unit is active across a set of stimuli. Population sparsity captures how selectively units respond to a given stimulus. To delineate how ReLU1 and our inhibitory modules contribute to activation sparsity, we computed activations from both Conv1 and its subsequent ReLU1 on the test set (*n* = 79 images).

#### Kurtosis

Kurtosis measures the tailedness of a distribution and can serve as an indicator of sparsity [Vinje and Gallant, 2000]. It is defined as:

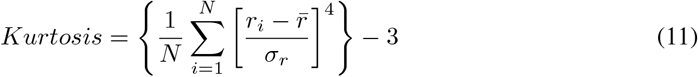

where *r*_*i*_ denotes individual sample values, 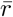 and *σ*_*r*_ are the mean and standard deviation of the sample, and *N* the number of elements. Higher values indicate a distribution with more extreme deviations.

#### Gini coefficient

The Gini coefficient [Gini, 1912], originally proposed to measure statistical dispersion, has been shown sufficient as a sparsity measure [Hurley and Rickard, 2009]. It is defined as:

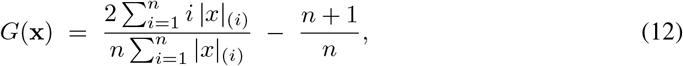

By convention, a uniform distribution’s Gini coefficient is 0, and this value approaches 1 as the distribution’s mass is concentrated in a few elements.

#### Hoyer sparsity

Hoyer sparsity [Hoyer, 2004] combines the *ℓ*_1_ and *ℓ*_2_ norms into a normalized score in [0, 1] and has been shown to be a cale-invariant sparsity measure [Hurley and Rickard, 2009]. It is defined as:

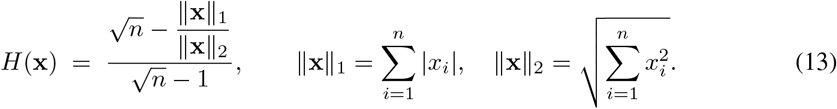

A dense vector of a uniform distribution has Hoyer sparsity of value 0, and 1 means this distribution only has one non-zero element.

#### Implementation details

All sparsity metrics were computed in Python. For weight sparsity, the Conv1 weight tensor (48, 2, 9, 9) was flattened per model seed. For activation sparsity, responses from Conv1 and ReLU1 had shape (*S, I, C, H, W*) = (10, 79, 48, 28, 56), where *S* denotes model seeds, *I* input images, *C* channels, and (*H, W*) spatial dimensions. This activation tensor was used to compute both *lifetime sparsity* and *population sparsity*. The former was computed for each feature channel. Specifically, we reshaped the tensor into (*S, C, I* · *H* · *W*), treating all activations of a given feature channel as a single sample for lifetime activation. Sparsity was computed along the last dimension, yielding a (*S, C*) matrix, which was then averaged across feature channels to obtain a (*S*,) vector. For *population sparsity*, all units’ activation for one stimulus are treated as the input for sparsity computation. In practice, we reshaped the tensor into (*S, I, C* · *H* · *W*) and calculated the sparsity on the last dimension, then further averaged across images.

#### Bootstrap and statistical analysis

For each model, all metrics (e.g., Test CC, Log-likelihood, *SI*) was evaluated across 10 random seeds on a held-out test set. Summary statistics were computed per model by taking the median across seeds. We estimated uncertainty via non-parametric bootstrapping: the 10 seed observations were resampled with replacement 10,00 times, the median was recomputed for each resample, and the 2.5th and 97.5th percentiles of the resulting distribution were taken as the bounds of the 95% confidence interval (CI). These intervals are reported as asymmetric error bars around the reported medians. Since the sample size is small (*n* = 10), bootstrap CIs should only be interpreted as descriptive.

To compare pairs of models (Baseline vs. Sub, baseline vs. Div and the corresponding pairs for BN model variants), we carried out paired analyses across the shared random seeds. For each comparison, we computed: (1) the median of the paired differences along with a bootstrap 95% CI, (2) the test statistic and p-value from a two-sided Wilcoxon signed-rank test, (3) the matched-pairs rank-biserial correlation as the effect size. We controlled the family-wise error rate using Holm-Bonferroni method at *α* = 0.05 for all comparisons from the same hypothesis. All levels of statistical significance are based on corrected p-values and are reported as * *p <* 0.05, ** *p <* 0.01.

## 3 Results

### 3.1 Inhibitory models and non-inhibitory models exhibit comparable predictive performance

After training each model, we tuned its hyperparameters with validation data and measured its predictive performance with test data (Figure 2a) using two complementary metrics: the Pearson correlation coefficient (CC) and the log-likelihood (LL). We refer to baseline models incorporated with either a subtractive or a divisive module as Sub or Div, baseline models with a normalization layer (BN, LN, or GN) as BN, LN, or GN. When both normalization and inhibition are used, they are referred as XN_Sub or XN_Div.

**Figure 2:**
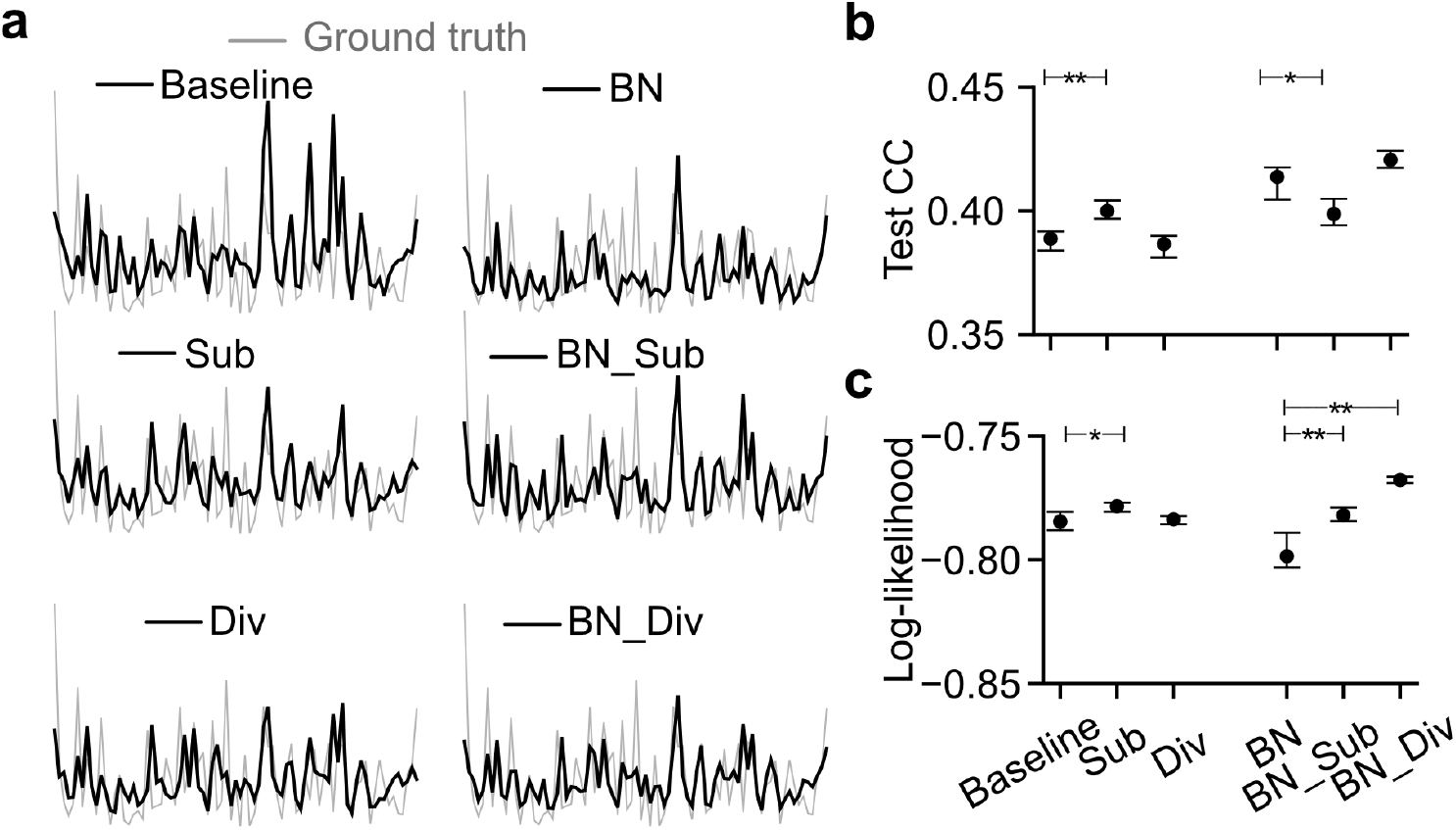
Predicting neuronal responses with inhibitory processing.. **(a)** Exemplary response traces from test data for one neuron (gray, recorded responses; black, predicted responses). **(b)** Predictive model performance measured by CC on test data (median across n = 10 random seeds per model). **(c)** Same as (b) but measured by log-likelihood. Error bars in (b) and (c) represent 2.5 and 97.5 percentiles obtained from bootstrapping.

When using CC as the performance metric (Figure 2b), we observed that the Sub model showed a modest improvement over the baseline (*p* = 0.008), whereas the Div model did not differ significantly from the baseline (*p* = 0.168). A similar pattern was observed on the second animal’s dataset (Suppl. Figures A.3). For BN variants, BN_Sub showed reduced CC relative to BN (*p* = 0.012), while BN and BN_Div did not differ significantly (*p* = 0.168). When evaluating performance using LL (Figure 2c), the baseline, Sub, and Div models exhibited a similar overall pattern to the CC-based evaluation. In contrast, for BN variants, both BN_Sub and BN_Div produced slightly higher LL values than BN despite the absence of corresponding improvements in CC.

For LN and GN variants, we observed that divisive models generally performed worse than their respective baselines, whereas subtractive models showed comparable or moderately improved performance (Suppl. Figures A.2).

Overall, these results indicate that incorporating subtractive or divisive operations as a proxy of inhibitory processing preserved predictive performance in some tested model families, while in some cases producing modest metric-dependent improvements in predicting neural responses to natural images.

### 3.2 Inhibition drives the emergence of biologically plausible representation

It has been reported that early stages along the visual pathway, including the retina and V1, feature antagonistic center-surround or localized oriented receptive fields (RFs) [Hubel and Wiesel, 1959, Chichilnisky, 2001, Qiu et al., 2023, Franke et al., 2022]. We assume that a network resembling biological visual systems exhibits internal representations similar to such feature selectivity.

We analyzed the filters of the first convolutional layer for each model trained by the first animal’s dataset (Figure 3a) and noticed that most filters of the baseline model and BN variants were quite noisy. Interestingly, Sub and Div models featured many filters that resembled Gaussian, Difference-of-Gaussian, or Gabor shapes (more spatial kernel examples in Suppl. Figure A.4). To quantify how much these learned filters are similar to the biological representation, we fit three parametric functions – a two-dimensional Gaussian, a Difference-of-Gaussian, and a Gabor function – to each filter. Next, we computed the respective *R*^2^ values as a measure of fit quality (Figure 3b). We used the maximum of the three *R*^2^ as a metric of biological plausibility.

**Figure 3:**
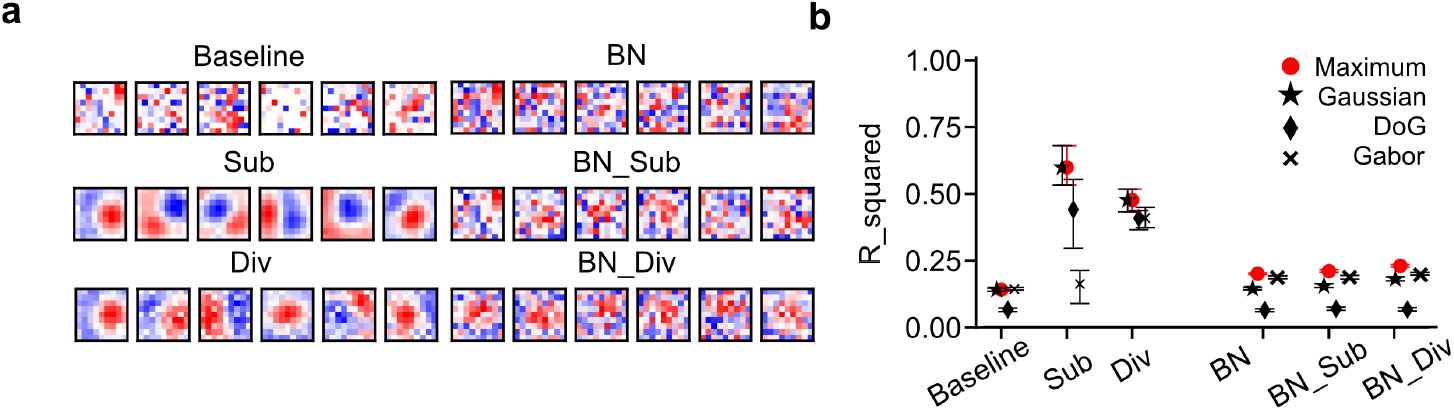
Neural representation for the trained models. **(a)** Exemplary filters of the first convolutional layer in the UV channel. **(b)** Median *R*^2^ of fitting a two-dimensional Gaussian (⋆), Difference-of-Gaussian (♦) or Gabor (×) function to convolutional kernels (a), and the maximum (red) of the three *R*^2^ values for each kernel. Error bars represent 2.5 and 97.5 percentiles obtained from bootstrapping.

We found that, compared to the Baseline, Sub and Div models achieved higher *R*^2^ values for fits with two-dimensional Gaussian and Difference-of-Gaussian, but not for Gabor (Figure 3b). The Div model showed high goodness-of-fit for all three functions. These observations indicate that our implementation of within-channel subtractive operation can drive the emergence of center-surround features, while the divisive module can facilitate the learning of oriented representations reminiscent of Gabor filters.

Additionally, we observed that BN variants had similar *R*^2^ values as the ones of the Baseline model (Figure 3b), suggesting that, although batch normalization improves predictive performance (cf. Figure 2b), it is not beneficial for the learning of biological features.

In summary, these results demonstrate that, compared to traditional models, networks using inhibition yield filters that resemble the feature selectivity observed in biological neural systems.

### 3.3 Inhibition benefits the learning of surround suppression

Visual systems exhibit nonlinear functions such as surround suppression [Cavanaugh et al., 2002a,b]. This phenomenon captures the reduction in a neuron’s response when a stimulus extends beyond the classical receptive field (CRF), inside which a stimulus directly drives a neuron’s responses. The area outside the CRF includes an inhibitory surround, which suppresses the neuron’s response when a stimulus extends beyond the CRF – a phenomenon known as surround suppression. Further outside this surround lies the extraclassical receptive field (eCRF), which can modulate the neural response without driving activity on its own. Surround suppression, arising from the inhibitory surround, has been observed across multiple stages of visual processing including retinal ganglion cells in frogs [Barlow, 1953], the lateral geniculate nucleus in monkeys [Krüger, 1977], and V1 in mice [Self et al., 2014], highlighting its broad functional role in visual processing. We examined whether adding inhibitory modules to a DNN would induce surround-suppression-like response patterns, analogous to those found in biological neurons.

To test this hypothesis, we conducted *in silico* experiments and computed the optimal Gabor filter for each neuron and for each model. After identifying the optimal Gabor, we generated a set of Gabor stimuli by varying the size and fed them to the trained model to get the corresponding model neurons’ responses. Three example cells’ modeled size-dependent responses are shown in Figure **??**a. For these three cells, both the Sub and the Div models exhibited surround suppression while the Baseline did not (More example neurons see Suppl. Figure A.8 and Suppl. Figure A.9 for mouse1 and mouse2, respectively). To evaluate this effect across neurons for each model, we computed the proportion of neurons with a suppression index (*SI*) greater than 0.1 (Figure 4b). We noticed that the Sub model displayed a higher fraction of surround-suppressed neurons than the Div model; still, the Div model had a higher fraction than the baseline model. The BN_Sub and BN_Div both had high fraction of neurons with *SI >* 0.1 higher the BN model.

**Figure 4:**
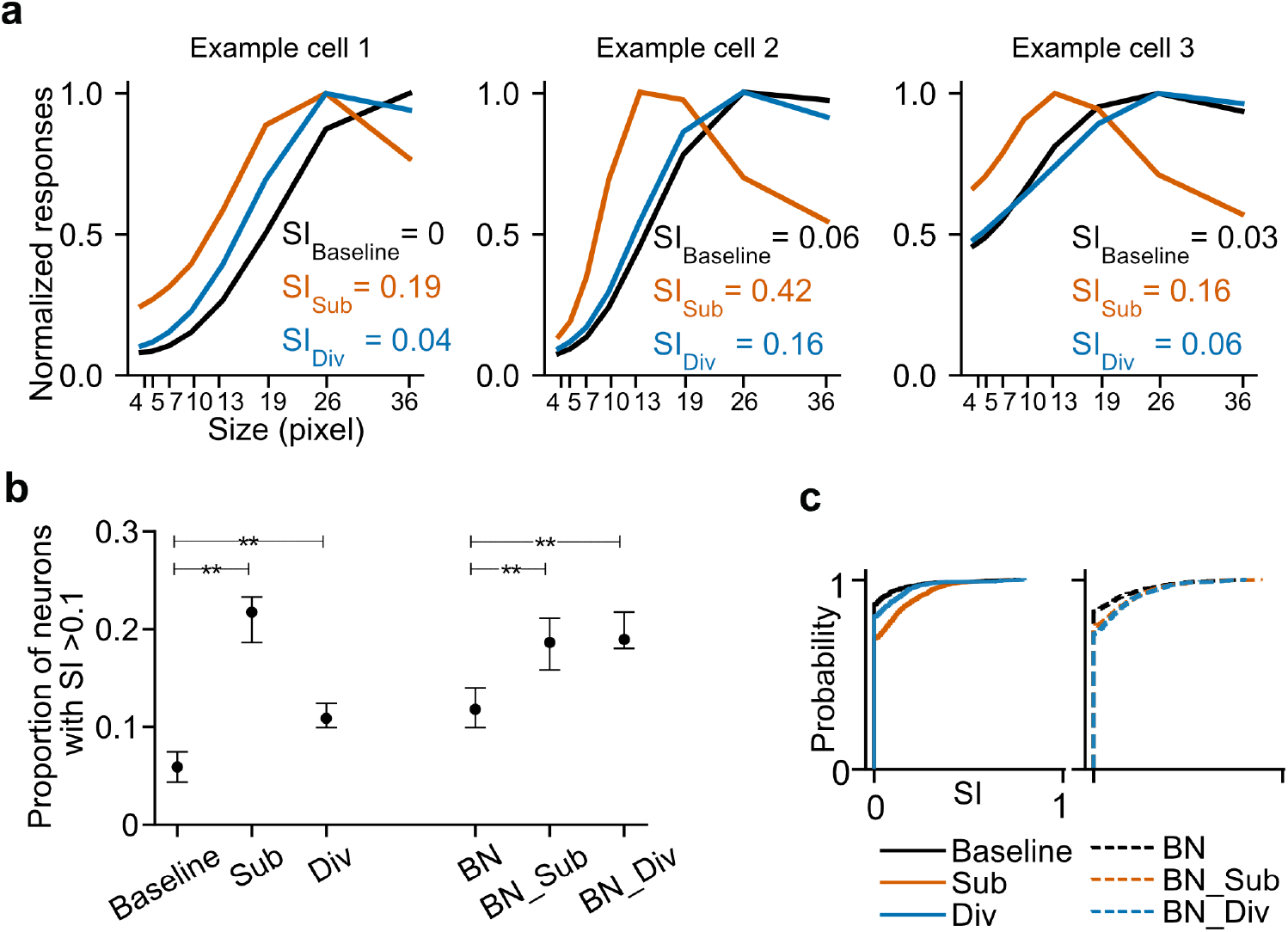
*In silico* experiments for testing surround suppression. **(a)** Tuning curves against size of Gabor stimulus for three exemplary cellx from three models without batch normalization (black, baseline; vermillion, subtraction; blue, division), with *SI* denoted. The subtraction and division model neurons’ response declined as the stimulus size increased, demonstrating the emulated surround suppression phenomenon. **(b)** Median fraction of neurons (n = 161) with surround-suppression index *SI >* 0.1. Error bars represent 2.5 and 97.5 percentiles obtained from bootstrapping. **(c)** Cumulative distribution function of *SI* for the 6 models.

To quantify how the *SI* values were distributed in each model, we plotted the cumulative distribution function of *SI* for all units per model. We observed that the Sub model exhibited a slow increase in its cumulative curve (Figure 4c), indicating a larger fraction of units with higher *SI* values relative to the Baseline. We also examined the distribution of *SI* in the LN and GN model variants (Suppl. Figure A.7) and found that the divisive module increased *SI* values in LN and GN, while the subtractive module produced higher *SI* only in LN.

Our results show that models incorporating inhibition – particularly the subtraction operation – successfully induced a high fraction of surround-suppressed neurons, suggesting that adding inhibitory interaction is an important step towards developing DNNs that more accurately emulate the complex response properties of biological systems.

### 3.4 Subtraction disrupts the emergence of cross-orientation inhibition

Similar to surround suppression, cross-orientation suppression is another well-studied nonlinear interaction in the visual cortex [Morrone et al., 1982]. It is characterized by a reduction in the response of an orientation-selective (OS) neuron to its preferred stimulus when an orthogonal stimulus is superimposed. While first described in cats [DeAngelis et al., 1992] and later reported in ferrets [Popović et al., 2018] and humans [Brouwer and Heeger, 2011], cross-orientation suppression has also been observed in mouse V1, primarily in the context of contrast normalization and broadly tuned suppression [Li et al., 2012, Liu et al., 2011]. We therefore examined whether incorporating inhibitory operations into our models drives the emergence of cross-orientation inhibition in our models.

To this end, for each neuron, we generated its orthogonal Gabor filter by rotating the corresponding optimal Gabor 90°. The orthogonal Gabors and their corresponding optimal Gabors were superimposed and formed a stimulus grid (Suppl. Figure A.10a). We then fed the overlaid stimulus to the trained model. For an exemplary neuron in the baseline model, we observed that cross-orientation inhibition was already present at the optimal Gabor contrast level of 7.1% (solid gray lines in Figure 5a). A similar trend was observed for this neuron in the Div and BN models. However, the Sub model showed the opposite trend: Neuronal responses increased with the increasing contrast of the orthogonal Gabors. The corresponding response matrices also showed that the BN neuron exhibited stronger cross-orientation inhibition than the Baseline, Sub, and Div neurons (Figure 5b).

**Figure 5:**
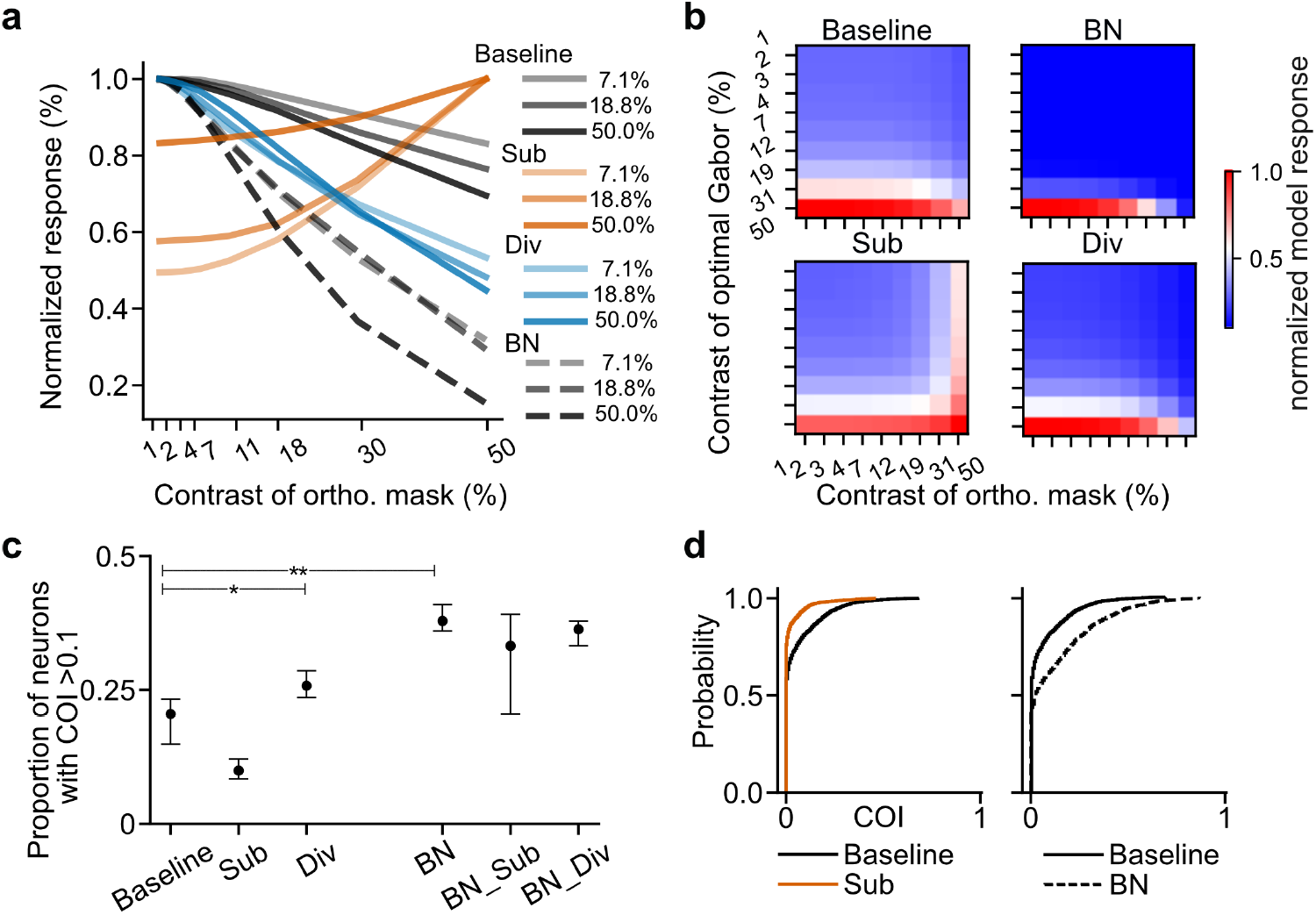
*In silico* experiments for testing cross-orientation inhibition. **(a)** Exemplary tuning curves against contrasts of orthogonal mask at different contrast levels of optimal Gabor for the Baseline (black), Sub (vermillion), Div (blue) and BN (dashed) models. **(b)** Example response matrix activated by a combination of optimal Gabor and orthogonal mask for the Baseline, Sub, Div, and BN models. **(c)** Median fraction of neurons (*n* = 161 per seed) with cross-orientation index *COI >* 0.1. Error bars represent 2.5 and 97.5 percentiles obtained from bootstrapping. **(d)** Cumulative distribution function of *COI* for the Baseline, Sub, and BN models.

To examine this effect at the single-neuron level, we computed the fraction of neurons with *COI >* 0.1 for each trained model (Figure 5c). Relative to the Baseline model, this fraction was higher in both the BN model (*p* = 0.010) and the Div model (*p* = 0.016), while the Sub model trended lower but did not reach significance after correction (*p* = 0.059). In addition, we also quantified the fraction of neurons with *COI >* 0.1 for LN, GN variants (Figure A.11). For the LN models, neither inhibitory module had an influence on the emergence of cross-orientation inhibition, whereas for the GN models the subtractive module significantly increased the fraction (*p* = 0.020). Furthermore, the cumulative distribution shows that the BN curve rises more slowly than that of the Baseline (Figure 5d), indicating a systematic shift toward higher *COI* values at the population level.

In summary, these results suggest that incorporating batch normalization improves learning cross-orientation inhibition while our implementation of within-channel subtraction impedes the learning of cross-orientation inhibition.

### 3.5 Inhibition changes weight sparsity and activation sparsity

Previous studies have demonstrated that incorporating subtractive or divisiv modules improves the sparsity of internal activations in DNNs trained for object recognition [Hasani et al., 2019, Miller et al., 2021]. Inspired by these results, we next examined whether our neural predictive models exhibited a similar effect, particularly in terms of connectivity sparsity and activation sparsity for the first convolutional layer (Conv1). We analyzed the distributions of both model weights and activations. Model activation is assessed in two standard forms: lifetime activation and population activation. We used kurtosis, Gini efficient, and Hoyer sparsity to quantify sparsity. Kurtosis captures the tailedness of a distribution, while the latter two metrics provide complementary but sufficient measures of distributional concentration and inequality [Rodieck, 1965]. By combining these three metrics, we obtained a more comprehensive characterization of the sparsity of distributions.

Histograms of model weight and activation revealed that the Sub model had higher fraction of weights near zero (Figure 6a), whereas units from Div model exhibited lower activation values (Figure 6b,c). We further characterized these distributions with the aforementioned three sparsity estimators. We observed that the subtractive module led to higher weight sparsity relative to the Baseline, whereas the divisive module produced the opposite pattern (Figure 6d,g,j). For activation sparsity, the effect diverged: the divisive modules increased both lifetime and population sparsity, while the subtractive module showed a weaker or inconsistent effects (Figure 6e,h,k for lifetime activation; f,i,l for population activation). To assess whether these effects persist after the first RuLU layer (ReLU1), we also examined the activations following ReLU1, confirming that similar effects remain for the Baseline, Sub, and Div models (gray box-plots in Figure 6d-i). Addtionally, we observed similar effects for BN variants. Compared to the BN model, the BN_Sub model had higher weight sparsity but lower lifetime and population activation sparsity, whereas the BN_Div showed the opposite effect. Furthermore, we tested how subtractive and divisive modules influence LN and GN models. We found that subtraction decreased the weight sparsity while division increased activation sparsity (Suppl. Figure A.12).

**Figure 6:**
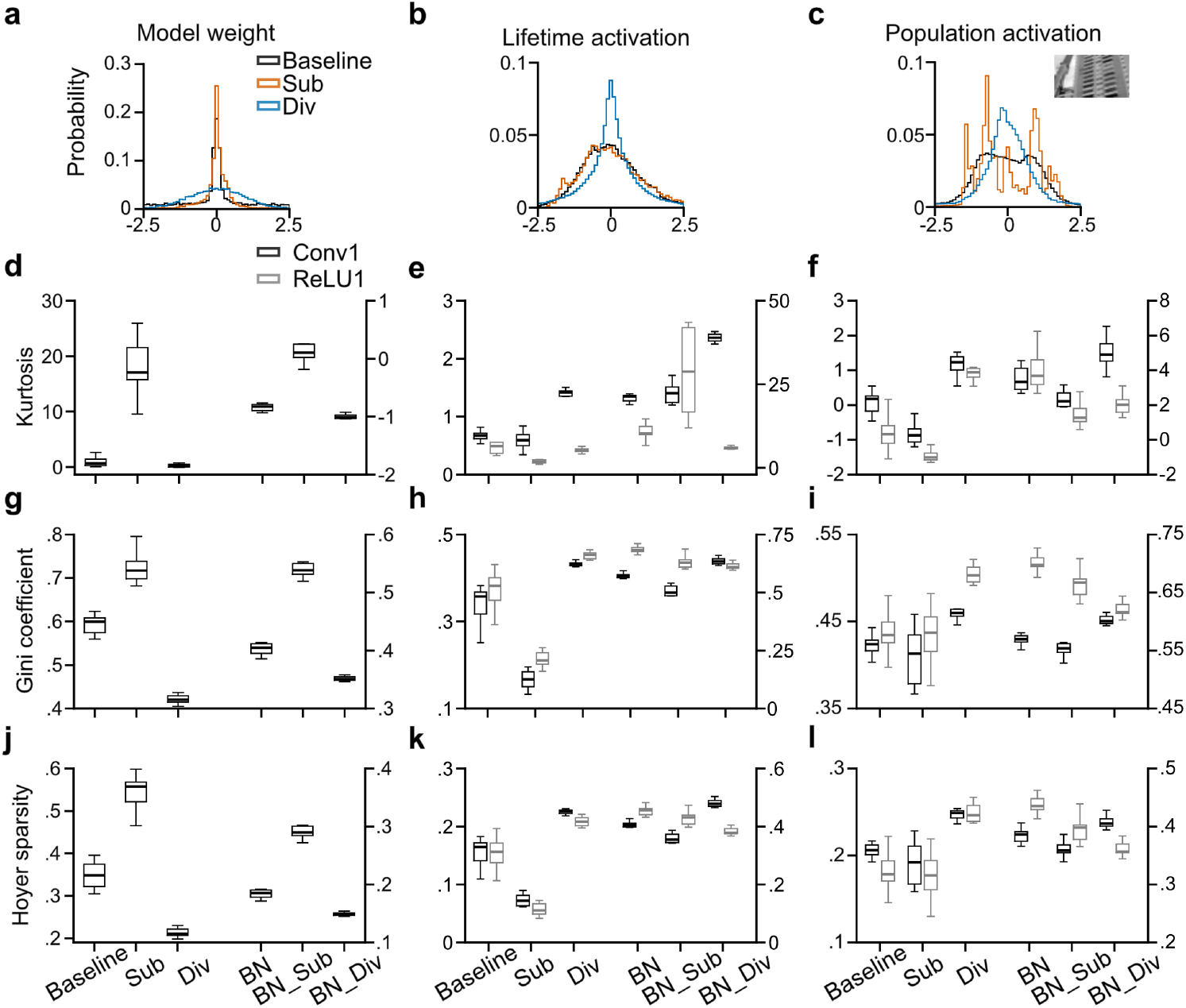
Weight and activation sparsity. **(a–c)** Z-transformed distributions for the Baseline, Sub, and Div models. (a). Z-scored model weights; (b). Z-scored lifetime activation distributions of one example unit; (c). Z-scored population activation distributions for the shown example test image. **(d–e)** Box plots of kurtosis values, n=10 random seeds for each model. (d). Kurtosis for model weights; (e). Kurtosis for lifetime activation before (black) and after (gray) ReLU; (f). Kurtosis for population activation before (black) and after (gray) ReLU. **(g–i)** Box plots of Gini coefficient values, n=10 random seeds for each model. (g). Gini coefficient for model weights; (h). Gini coefficient for lifetime activation before (black) and after (gray) ReLU; (i). Gini coefficient for population activation before (black) and after (gray) ReLU. **(j–l)** Box plots of Hoyer sparsity values. (j). Hoyer sparsity for model weights; (k). Hoyer sparsity for lifetime activation before (black) and after (gray) ReLU; (l). Hoyer sparsity for population activation before (black) and after (gray) ReLU. The significance test results for Baseline vs. Sub and Baseline vs. Div are reported in Table 1.

In summary, our results suggest that the subtractive operation increases connectivity sparsity but decreases activation sparsity. In contrast, the divisive operation improves the activation sparsity while reducing connectivity sparsity.

**Table 1:** Tests results for paired non-parametric comparisons of three sparsity estimators across *n* = 10 random seeds in Figure 6).

| Sparsity | Estimator | Comparison | Median $\Delta$ | 95% CI | $r_{rb}$ | $p_{corr}$ |
| --- | --- | --- | --- | --- | --- | --- |
| Weight | Kurtosis | vs. Sub | -16.11 | [-21.9, -13.9] | -1.00 | .0039* |
|  |  | vs. Div | 0.55 | [-0.08, 1.75] | 0.67 | .064 |
|  | Gini | vs. Sub | -0.13 | [-0.16, -0.10] | -1.00 | .0039* |
|  |  | vs. Div | 0.17 | [0.15, 0.19] | 1.00 | .0039* |
|  | Hoyer | vs. Sub | -0.21 | [-0.23, -0.19] | -1.00 | .0039* |
|  |  | vs. Div | 0.14 | [0.11, 0.16] | 1.00 | .0039* |
| Lifetime | Kurtosis | vs. Sub | 0.05 | [-0.00, 0.19] | 0.67 | .0644 |
|  |  | vs. Div | -0.72 | [-0.78, -0.62] | -1.00 | .0039* |
|  | Gini | vs. Sub | 0.18 | [0.12, 0.21] | 1.00 | .0039* |
|  |  | vs. Div | -0.07 | [-0.12, -0.06] | -1.00 | .0039* |
|  | Hoyer | vs. Sub | 0.09 | [0.05, 0.11] | 1.00 | .0039* |
|  |  | vs. Div | -0.06 | [-0.08, -0.05] | -1.00 | .0039* |
| Population | Kurtosis | vs. Sub | 0.92 | [0.47, 1.23] | 1.00 | .0039* |
|  |  | vs. Div | -1.09 | [-1.50, -0.76] | -1.00 | .0039* |
|  | Gini | vs. Sub | 0.02 | [-0.02, 0.04] | 0.345 | .3750 |
|  |  | vs. Div | -0.04 | [-0.05, -0.02] | -1.00 | .0039* |
|  | Hoyer | vs. Sub | 0.02 | [-0.01, 0.03] | 0.564 | .1308 |
|  |  | vs. Div | -0.04 | [-0.05, -0.03] | -1.00 | .0039* |

## 4 Discussion

In this study, we examined the effects of incorporating inhibitory processing as an inductive bias into a DNN to identify visual response functions. By implementing subtractive and divisive operations, we demonstrated that, compared to our baseline model, networks that incorporate inhibition obtained comparable predictive performance while being encouraged to learn biologically plausible representations. Our *in silico* experiments suggest that such operations drive the emergence of surround suppression but not cross-orientation inhibition. Additionally, we observed that the subtraction and division operations had reversed effects on the sparsity of our model activations. These results suggest the importance of considering inhibition for studying nonlinear and high-dimensional information processing in biological neural networks.

While relatively little effort has been devoted to incorporating inhibition for neural prediction (but see, e.g., [Burg et al., 2021] for an exception), extensive work has explored the use of DoG or DN in image recognition tasks. For example, the use of DoG in deep models has been shown to improve classification accuracy and generalize well to objects under different lighting conditions and occlusions [Hasani et al., 2019]. Pogoncheff et al. [2023] trained models on ImageNet and fine-tuned them on Tiny-ImageNet to evaluate their alignment with V1. They found that DoG modules had a marginal effect on predicting neural responses, yet surprisingly impaired the model’s property of surround modulation. In parallel, implementing local DN operations by inducing competition among feature channels has proven to be beneficial for image recognition. It increases activation sparseness and sharpens the activation map of early convolutional layers [Miller et al., 2021, Cirincione et al., 2022, Zhi et al., 2025, Pan et al., 2021, Ren et al., 2016]. Adding DN to a front-end convolutional layer – mimicking computations in the V1 – also improves performance in classifying corrupted images and enhances alignment with V1 properties such as surround suppression [Cirincione et al., 2022].

Note that our current implementation of inhibitory processing accounts for the interactions among neurons with nearby receptive fields [Hasani et al., 2019, Aqil et al., 2021], but does not model inhibition across feature channels [Miller et al., 2021]. This deviates from the canonical definition of divisive normalization, which directly implements interaction across feature channels. However, our tests with BN variants also demonstrate the possibility of modeling cross-orientation inhibition via within-channel rescaling. Future work could explore how different forms of inhibition and how they interact with normalization modules could influence the model’s ability to identify neuronal response functions.

We tested the effects of subtractive and divisive operations on surround suppression and cross-orientation inhibition. In the future, it would be interesting to examine the influence on visual components such as texture, color, shape, and motion [Yu et al., 2022, Jarvers and Neumann, 2024, Poleg-Polsky, 2025], which might be related to the presented stimulus types such as natural images, natural video, noise, and random dot kinematograms [Qiu et al., 2021, Turishcheva et al., 2024b,a]. In this case, a dynamic model would be used, which implements inhibitory computation in the temporal domain, offering an opportunity to test how visual systems adapt to changing environments [Carandini and Heeger, 2012, Młynarski and Hermundstad, 2021, Smeds et al., 2019, Qiu and Euler, 2019, Wang et al., 2025]. Additionally, while we focused on the kernels of the first convolutional layers, further analysis of other hidden representations would be insightful, given the hierarchical processing in the neural system.

Importantly, a closed-loop biological experiment design is required to provide the necessary support for our simulation results, in which static natural images and artificial stimuli, such as optimal Gabors, are deployed. We note that some of the findings, including the learned kernels and the sparsity of weights, may be challenging to examine. In general, our study suggests the non-triviality of considering functional features of the brain for building digital twin models.

## 5 Acknowledgments

We thank Fabian Sinz for helpful discussions. This work was supported by the German Research Foundation (DFG; CRC 1233 “Robust Vision: Inference Principles and Neural Mechanisms”, project number 276693517 to TE). AST acknowledges the support of the National Science Foundation and DoD OUSD (R&E) under Cooperative Agreement DBI-2229929 (The NSF AI Institute for Artificial and Natural Intelligence). This work was also supported by the National Institutes of Mental Health grant UM1MH130981, the National Institute of Neurological Disorders and Stroke (NINDS) grant R01 NS113890, the Intelligence Advanced Research Projects Activity (IARPA) via Department of Interior/Interior Business Center (DoI/IBC) contract number D16PC00003 and the James Fickel Enigma Project Fund. The funders had no role in study design, data collection and analysis, decision to publish, or preparation of the manuscript.

## 7 Data and Code Availability

The data and code will be available upon publication.

## 8 Author contributions

**Y. D**.: conceptualization, methodology, software, validation, formal analysis, investigation, data curation, writing (original draft), visualization **Z. D**.: validation, investigation, data curation, formal analysis, writing (review and editing) **J. F**.: data curation, methodology **O. J**.: formal analysis, writing (review and editing) **A. T**.: supervision, funding acquisition **T. E**.: resources, writing (review and editing), supervision, funding acquisition **Y. Q**.: conceptualization, methodology, software, formal analysis, investigation, writing (original draft), visualization, supervision, project administration

