## Supplemental_Figure_A9 for "Inhibition benefits neural system identification"

### A.9 10 example neurons exhibiting surround suppression in mouse 2 (n=91 cells) modeled by baseline, Sub, and Div models across 10 seeds

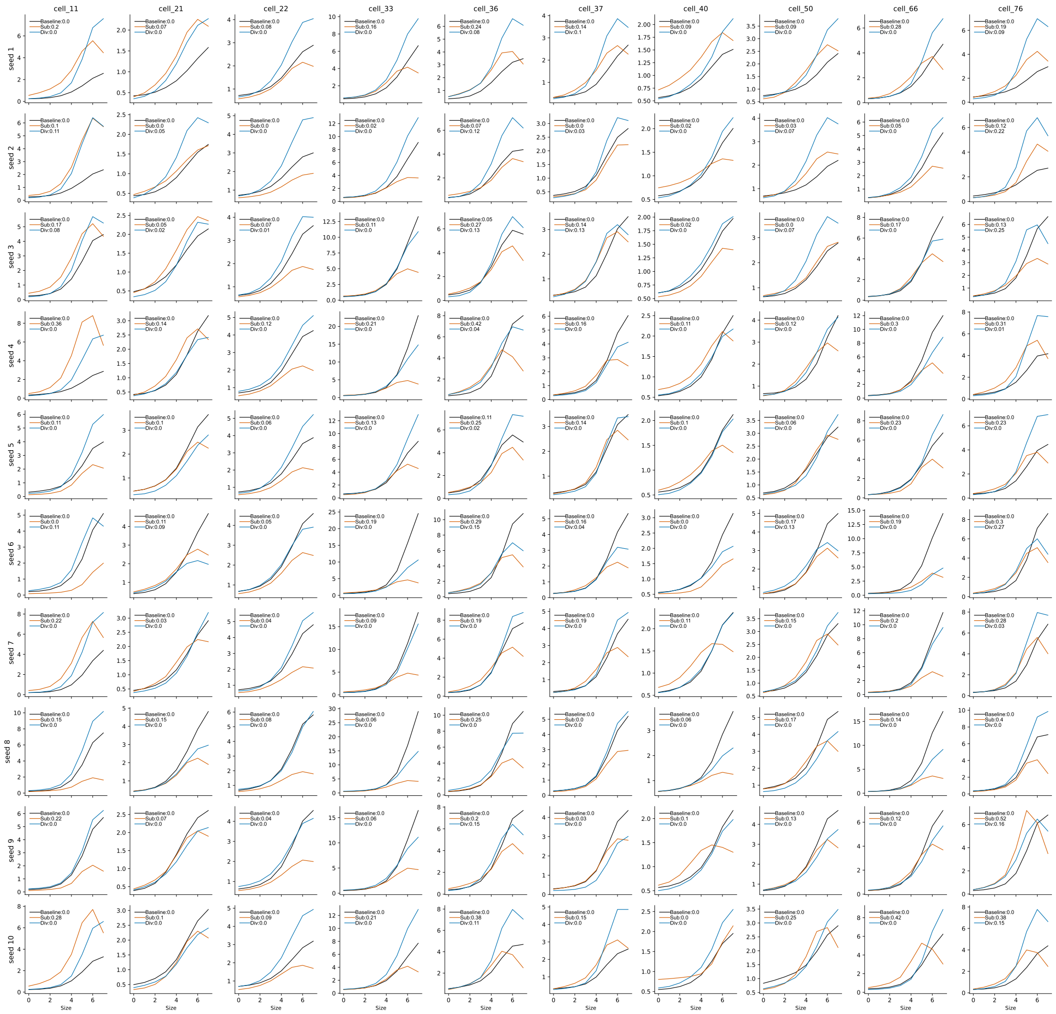
