## Supplemental_Figure_A1 for "Inhibition benefits neural system identification"

### A Appendix / supplemental material

#### A.1 Model architectures for baseline, BN, LN, GN and their inhibitory model variants

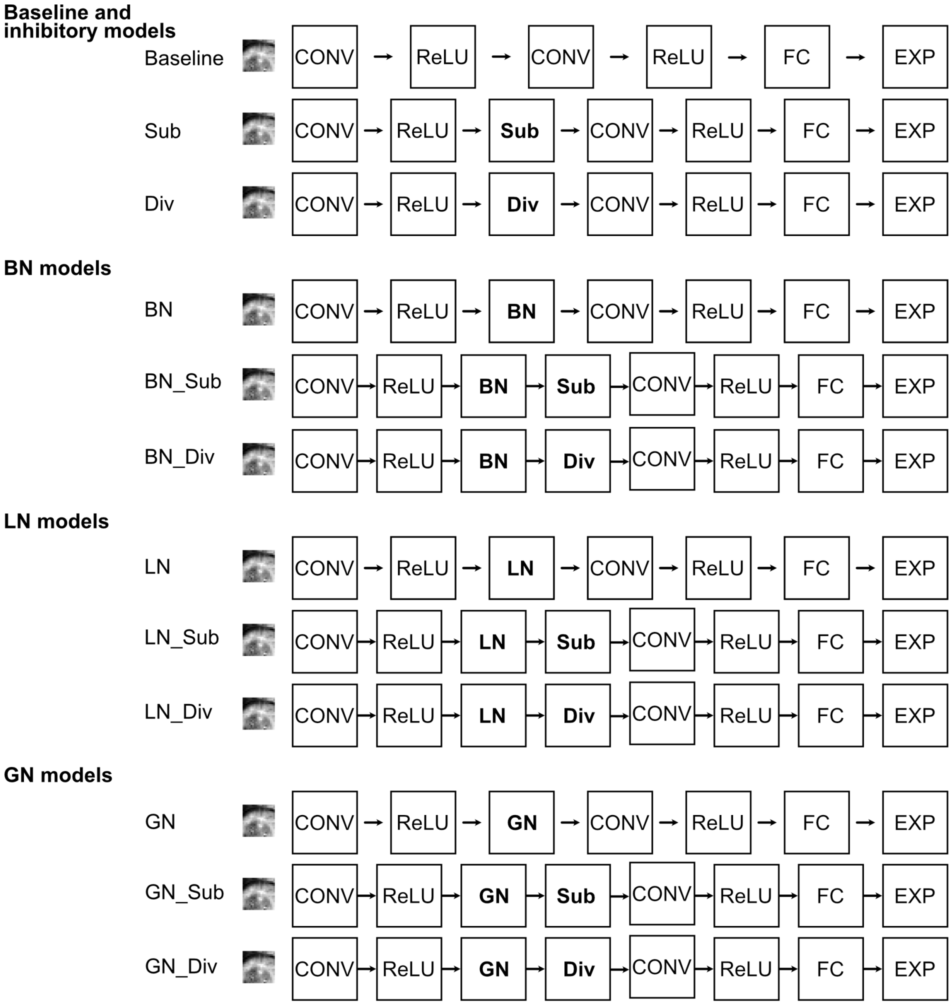


**Figure S 1** Model structures for baseline, BN, LN, GN, together with their respective inhibitory model variants. The input image has dimensions of $2 \times 36 \times 64$ (channels × height × width). The outputs of the first convolutional layer, first ReLU, inhibitory module, and normlization layer all have dimensions $48 \times 28 \times 56$.The outputs of the second convolutional and its subsequent ReLU have dimensions $48 \times 22 \times 50$.
