## Supplemental_Figure_A2 for "Inhibition benefits neural system identification"

### A.2 Predicting neuronal responses with inhibitory processing (Mouse 1, n=161 cells)

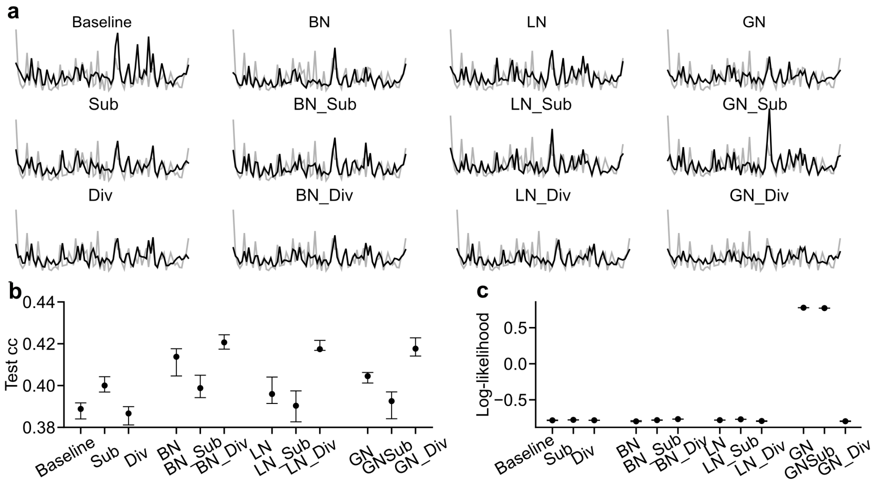


**Figure S 2.1 Predicting neuronal responses with inhibitory processing. a.** Exemplary response traces from test data for a neuron (gray, recorded responses; black, predicted responses from different models). **b.** Performance of the predictive model measured by the Pearson correlation coefficient (CC) in the test data (median across n = 10 random seeds per model). **c.** Same as in (b), measured by logarithmic likelihood. The error bars in (b) and (c) represent 2.5th and 97.5th percentiles obtained from bootstrapping.
