## Supplemental_Figure_A5 for "Inhibition benefits neural system identification"

### A.5 Exemplary spatial filters of the first convolutional layer of baseline, Sub, Div, BN, BN_Sub, BN_Div models, trained on mouse 2

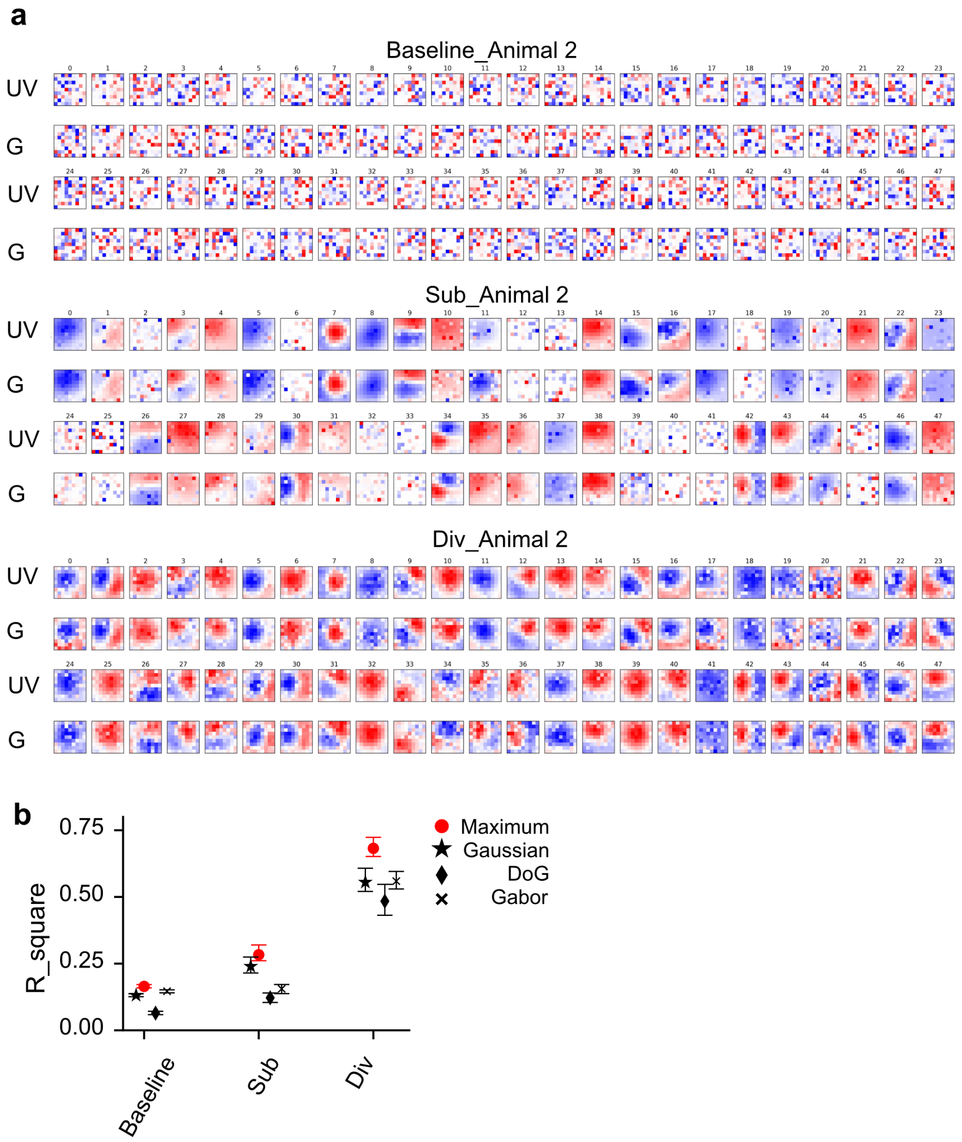


**Figure S 3.2 a.** 96 spatial filters of the first convolutional layer, from a randomly chosen model seed. The kernel size is $9 \times 9$ **b.** Median $R^2$ of fitting a two-dimensional Gaussian (★ ), Difference-of-Gaussian (♦) or Gabor ($\times$) function to convolutional kernels (a), and the maximum (red) of the three $R^2$ values for each kernel. Error bars represent 2.5 and 97.5 percentiles obtained from bootstrapping.
