## Supplemental_Figure_A6 for "Inhibition benefits neural system identification"

### A.6 Depiction of six parameters altered in the process of identifying the optimal Gabor

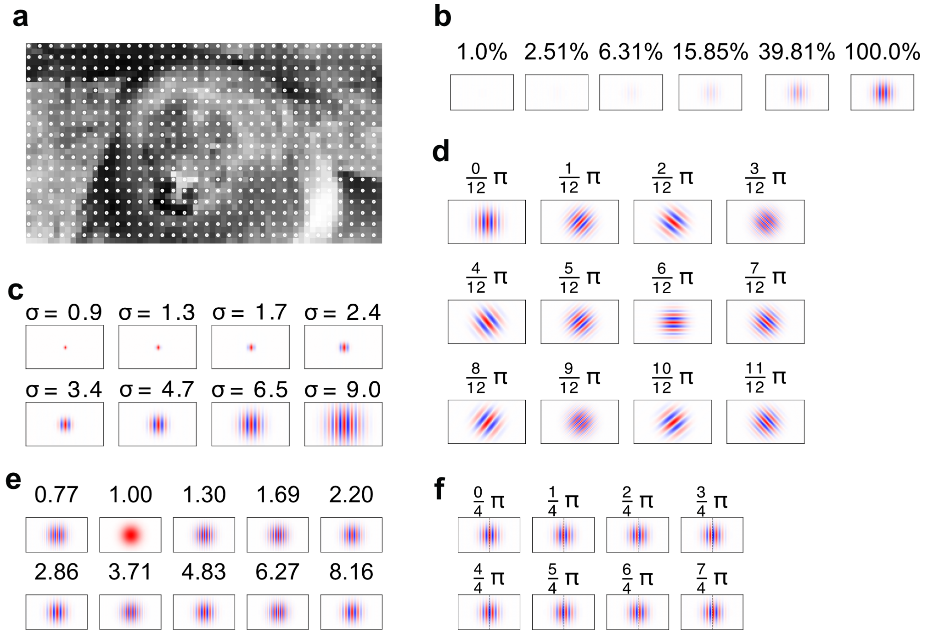


**Figure S 4.1 Depiction of six parameters altered in the process of identifying the optimal Gabor.** **a.** The 18 x 32 white dots scattered on an exemplary image (36 x 64) represent all 576 possible center positions of an optimal Gabors. Each white dot is two pixels away from its four-connected neighbors. **b.** Contrast. **c.** Standard deviation of the Gaussian envelope. **d.** Orientation. **e.** Spatial frequency. **f.** Phases.
