## Supplemental_Figure_A7 for "Inhibition benefits neural system identification"

### A.7 Overview of how surround suppression is modeled by the baseline, BN, LN, GN, and their inhibitory model variants in two mice

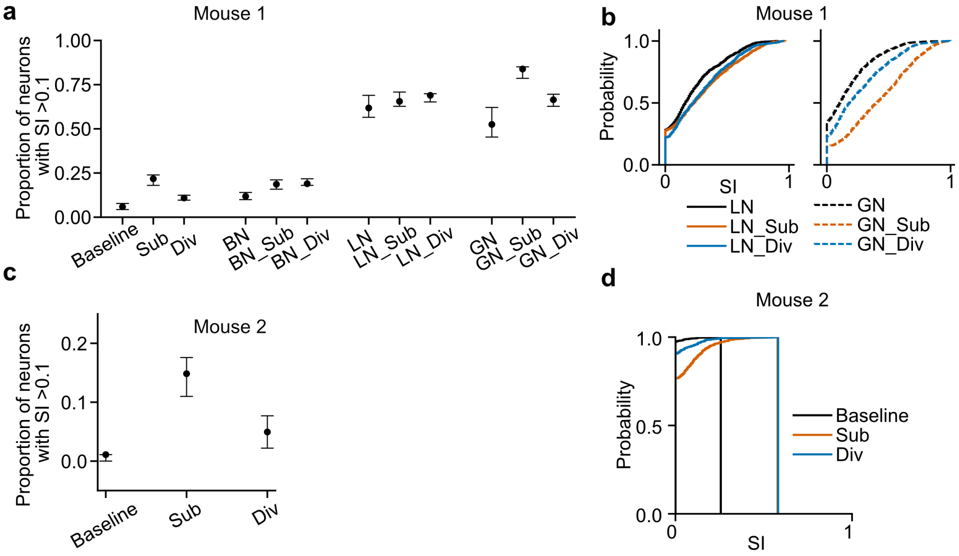


**Figure S 4.3** **a** Median proportion of neurons $(n = 161)$ per model seed, whose $SI > 0.1$ for the baseline, BN, LN, and GN and their inhibitory model variants in mouse 1. **b** Cumulative distribution function of $SI$ for the LN variants and GN variants in mouse 1. **c** Median proportion of neurons $(n = 91)$ per model seed, whose $SI > 0.1$ for the baseline, Sub, and Div in mouse 2. **d** Cumulative distribution function of $SI$ for the baseline, Sub, and Div in mouse 2. The error bars represent 2.5 and 97.5 percentiles obtained from bootstrapping.
