## Supplemental_Figure_A8 for "Inhibition benefits neural system identification"

### A.8 10 example neurons exhibiting surround suppression in mouse 1 (n=161 cells) modeled by baseline, Sub, and Div models across 10 seeds

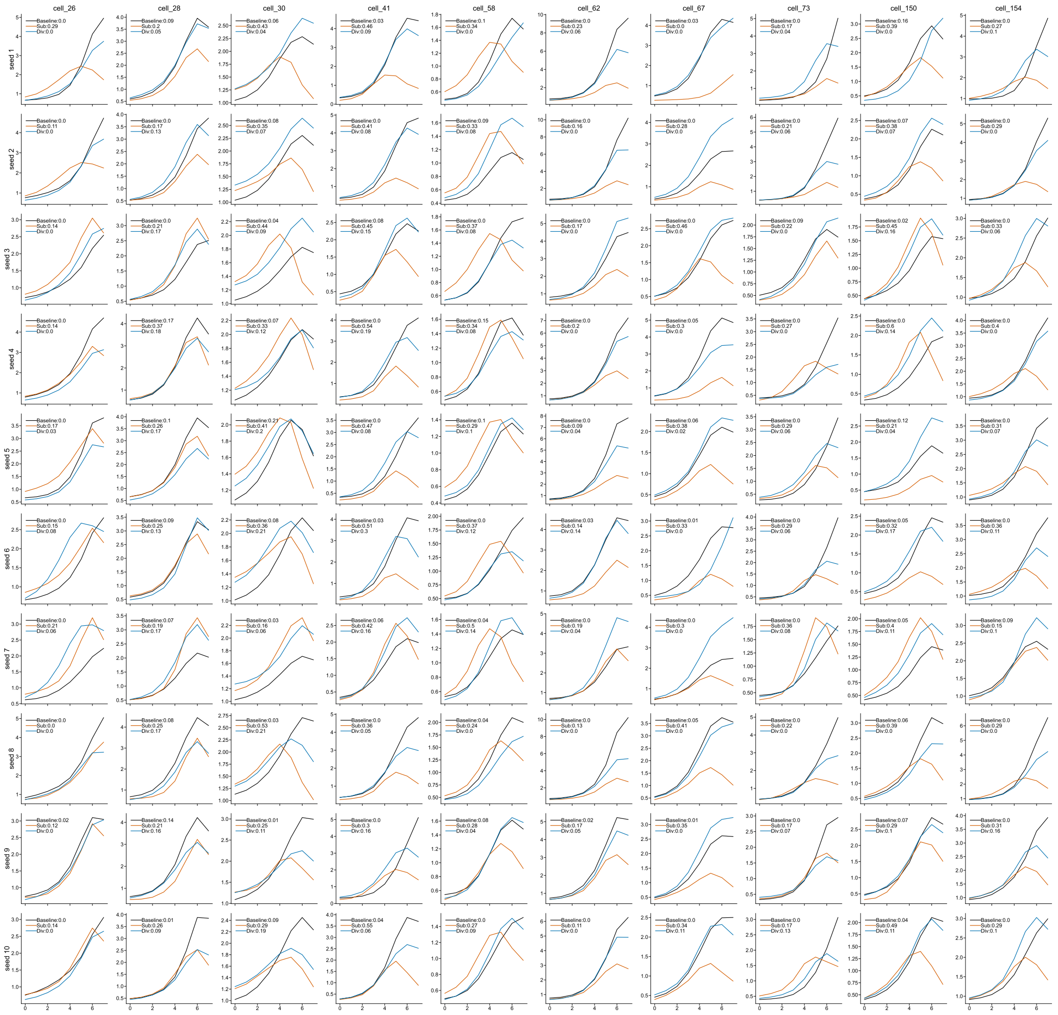
