## Supplemental_Figure_A10 for "Inhibition benefits neural system identification"

### A.10 Example stimulus and response grids for one example neuron in BN model

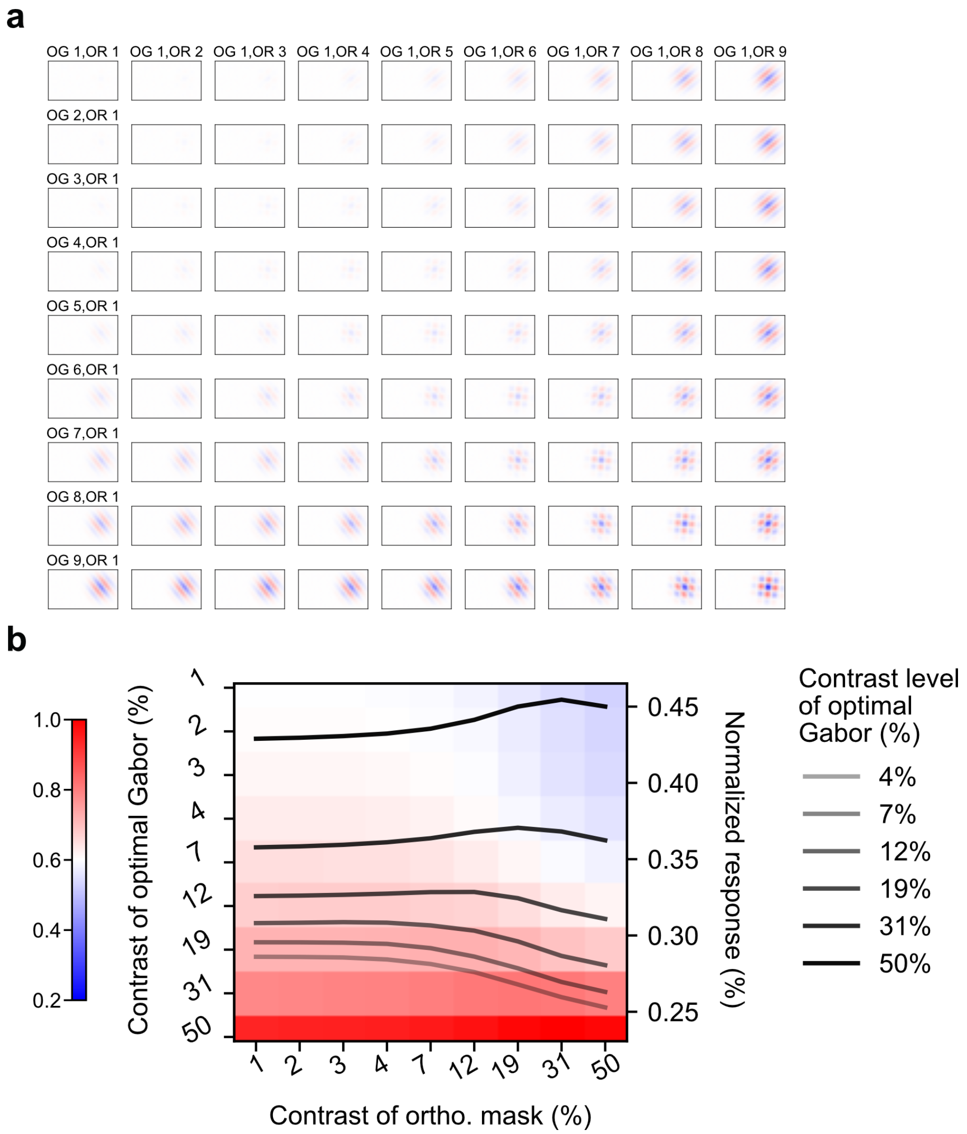


**Figure S 5.1 Example stimulus and response grids for one example neuron in BN model. a.** Gabors are arranged so that each row shares the same optimal contrast, while each column shares the same orthogonal contrast. The Gabor positioned on the top-left exhibits the lowest optimal and orthogonal contrasts, whereas the one at the bottom-right displays the highest values for both. **b.** One example BN cell's response matrix to the stimulus grid in (a). 6 lines depict how the model neuron's response changes as the superimposed orthogonal Gabor's contrast increases when the optimal Gabor's contrast were kept constant.
