## Supplemental_Figure_A11 for "Inhibition benefits neural system identification"

### A.11 In silico experiments for testing cross-orientation inhibition for all 12 models in mouse 1

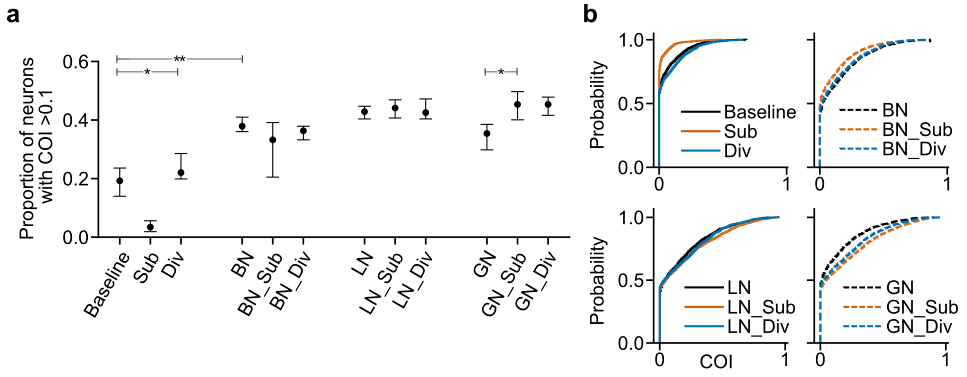


**Figure S 5.2 a.** Median proportion of neurons $(n = 161)$ with COI $> 0.1$ for all models. Error bars represent 2.5 and 97.5 percentiles obtained from bootstrapping. **b.** Cumulative distribution function of COI for all the models.
