## Supplemental_Figure_A12 for "Inhibition benefits neural system identification"

### A.12 Weight and activation sparsity across all models for mouse 1

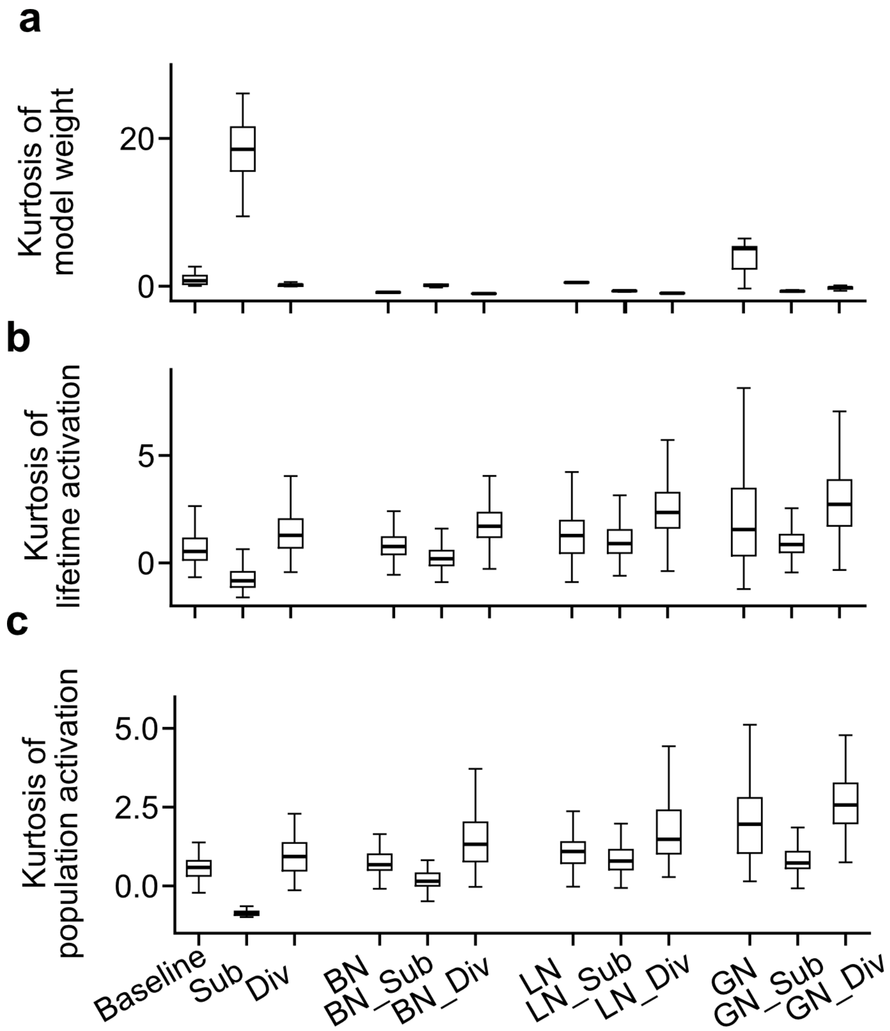


**Figure S 6.2 Weight and activation sparsity. a.** Box plots of weight kurtosis for all 12 trained models. **b.** Box plots of lifetime kurtosis for all 12 trained models. **c.** Box plots of population kurtosis for all 12 trained models.
